# AAV-mediated gene therapy for SLC13A5 Citrate Transporter Disorder rescues epileptic and metabolic phenotypes

**DOI:** 10.1101/2025.07.03.663044

**Authors:** Lauren E. Bailey, Morgan K. Schackmuth, Raegan M. Adams, Irvin T. Garza, Krishanna Knight, Sydni K. Holmes, Meghan M. Eller, MinJae Lee, Rachel M. Bailey

## Abstract

SLC13A5 citrate transporter disorder is a rare epileptic encephalopathy caused by loss of function pathogenic variants in the *SLC13A5* gene. Loss of sodium/citrate cotransporter (NaCT) function causes a severe early life epilepsy resulting in life-long developmental disabilities and increased extracellular citrate. Current antiseizure medications may reduce seizure frequency, yet more targeted treatments are needed to address the epileptic and neurodevelopmental SLC13A5 phenotype. We performed preclinical studies in SLC13A5 deficient mice evaluating phenotype rescue with adeno-associated virus (AAV) vector carrying a functional copy of the human *SLC13A5* gene (AAV9/SLC13A5). Cerebrospinal fluid-delivery of AAV9/SLC13A5 decreased extracellular citrate levels, normalized electrophysiologic and sleep architecture abnormalities, and restored resistance to chemically induced seizures and death. Treatment benefits were achieved with administration during early brain development and in young adult mice, supporting a broad therapeutic window for this disorder. Comparison of delivery routes in young adult KO mice showed that higher brain targeting achieved with intra-cisterna magna delivery resulted in greater treatment benefit as compared to intrathecal lumbar puncture delivery. Together, these results support further development of AAV9/SLC13A5 for treating SLC13A5 citrate transporter disorder.

**Graphical Abstract:** 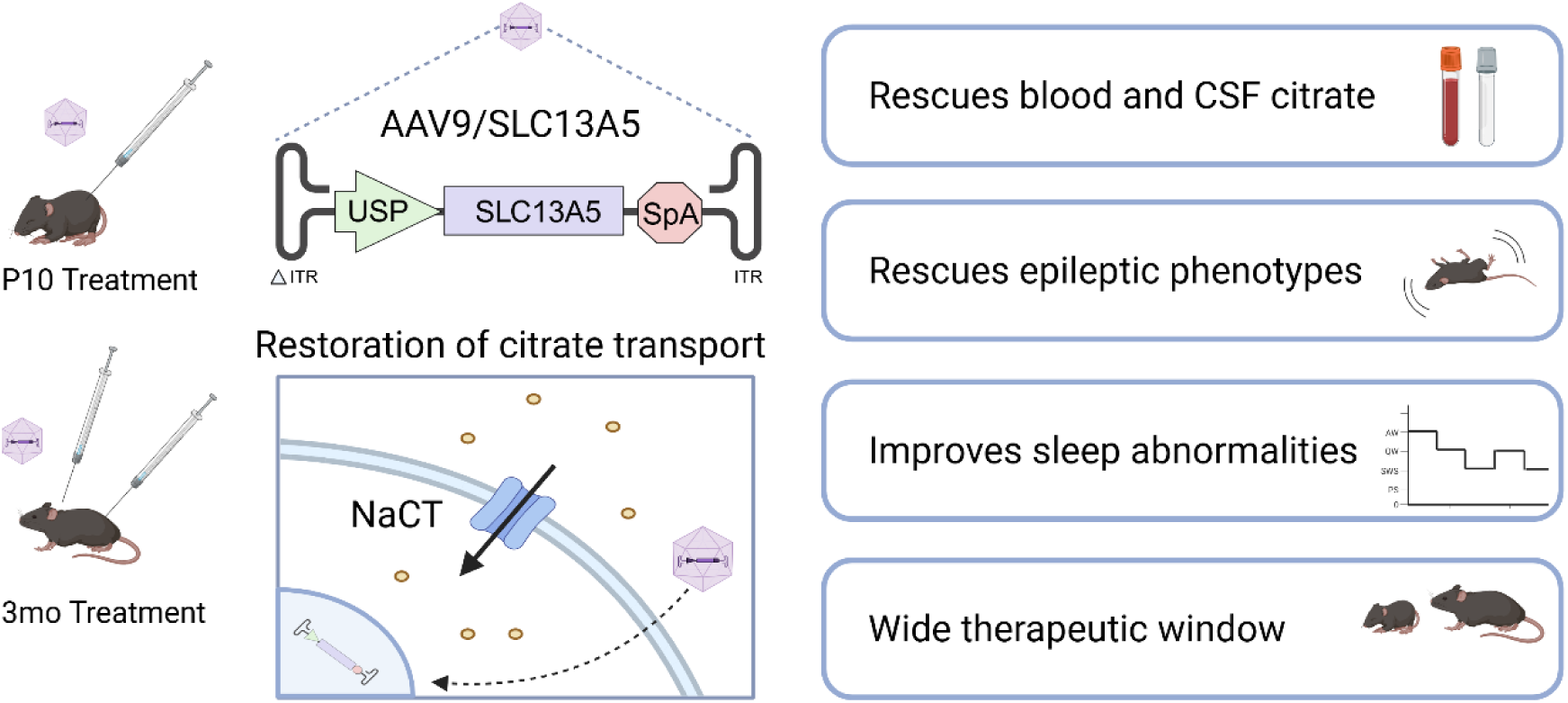

## INTRODUCTION

SLC13A5 citrate transporter disorder (also known as SLC13A5 Epilepsy, SLC13A5 Deficiency, or Developmental Epileptic Enchephalopathy-25 [DEE25]) is a rare autosomal recessive epileptic encephalopathy with seizure onset in the first day of life along with persistent intellectual and motor disabilities (1). Patients have multi-focal epilepsy, cognitive and sleep abnormalities (2). The disease is caused by bi-allelic loss-of function mutations in the *SLC13A5* gene (3). There are currently only symptomatic treatments and a curative therapy for SLC13A5 deficiency is needed. Seizures are often pharmacoresistant and children have succumbed to seizure complications (4). The degree of neurodevelopmental disability and seizure-burden vary, but most patients are severely affected, requiring 24/7 lifelong care (5). A therapy that addresses the underlying cause of the disorder would have a profound impact on the patients and their families’ wellbeing and decrease healthcare costs.

The *SLC13A5* gene codes for a sodium dependent citrate transporter (NaCT) that has the highest expression in the liver, followed by brain, testis, kidney, and bones. In brain, NaCT is found in the cell membrane of neurons, astrocytes, and other cell types. Citrate plays a critical role in cellular energy metabolism, neurotransmitter synthesis, and lipid metabolism. To date, all tested mutations result in no or a greatly reduced amount of citrate transport (6). SLC13A5 patients show distinctive elevations of citrate in the plasma and cerebrospinal fluid (CSF) (7-9). Diminished intracellular citrate and/or elevated extracelluar citrate may be the underlying cause of the frequent seizures and developmental disabilities in patients.

The *Slc13a5* knockout (KO) mouse lacking NaCT shows increased neuronal excitability and propensity for epileptic seizures, abnormal citrate levels in cerebrospinal fluid (CSF) and brain tissue, and impaired bone and tooth development (8, 10-12). Recently, we reported that young adult *Slc13a5* KO mice have sleep deficits, similar to the sleep deficits observed in patients, which coincided with abnormal EEG power spectrum (2). While *Slc13a5* KO mice model salient aspects of SLC13A5 citrate transporter disorder that can be utilized for therapeutic development they lack overt behavioral, motor or cognitive impairments that are seen in SLC13A5 patients (6).

Gene therapy is a precision medicine where functional genetic material is delivered to cells that has the potential to address the underlying cause of a disorder rather than managing symptoms. Adeno-associated virus (AAV) vectors have efficient gene transfer, broad serotype-dependent tropism, low risk of insertional mutagenesis, and long-term transgene expression in transduced cells following a single injection (13). AAV-based gene therapies show great promise in treating neurological disorders and there are currently six FDA approved AAV gene therapies using different serotypes (14-17). AAV serotype 9 (AAV9) is most frequently used for CNS brain delivery due to its ability to achieve broad transgene transduction across the CNS of rodents and large animals and injection into CSF transduces peripheral tissues in addition to the CNS (18, 19). Importantly, CSF AAV9 gene therapy is currently being utilized in active clinical trials for pediatric neurological disorders, such as giant axonal neuropathy, Rett Syndrome, and spastic paraplegia 50 (20-22).While AAV vectors are small, with a packaging size <5 kilobases, the coding sequence of the *SLC13A5* gene is within this limit, making gene therapy an amenable approach (23). Here we report the development of a self-complementary AAV9 vector delivering optimized *SLC13A5* (AAV9/SLC13A5) to treat SLC13A5 citrate transporter disorder, the design of which can be readily translated for clinical use. Preclinical efficacy studies were performed using CSF delivery in both pups and young adult *Slc13a5* KO mice. Following intrathecal lumbar puncture (IT) administration of AAV9/SLC13A5 in *Slc13a5* KO pups, blood and CSF citrate levels were decreased, brain activity and sleep abnormalities were normalized, epileptic discharges were decreased, and the sensitivity to seizure onset and severity were attenuated. In young adult mice, IT and intra-cisterna magna (ICM) administration of AAV9/SLC13A5 were directly compared and similar to adolescent mice, AAV9/SLC13A5 treatment decreased blood citrate levels, normalized EEG activity, improved sleep architecture, increased resistance to seizures, and protected against seizure-induced deaths. A greater treatment benefit in adult KO mice was achieved with ICM delivery, which also resulted in greater AAV9/SLC13A5 gene transfer in the adult brain. Importantly, AAV9/SLC13A5 administration in both pups and adult mice was safe and well tolerated.

## RESULTS

### Design and functional testing of a gene vector expressing NaCT

For downstream human application, we developed the AAV/UsP-SLC13A5 vector consisting of a ubiquitously expressed minimal synthetic JeT plus intron promoter (UsP) (24), driving the expression of codon-optimized human *SLC13A5* cDNA that has a synthetic polyA tail (Figure 1A) (25). Given AAV packaging constraints (26, 27), use of the short UsP promoter allows self-complementary packaging of the 1.7 kb SLC13A5 coding sequence, which is predicted to stably transduce at least 10-fold more cells than single stranded AAV (26, 27). The gene insert is bounded by the AAV2 ITRs, where one terminal repeat has the wildtype 144 nt sequence and the other ITR (ΔITR) is mutated to delete the AAV DNA resolution site and D sequence to direct preferential replication and packing of scAAV DNA sequences (28). Since SLC13A5 is expressed in both the brain (neurons and glia) and liver, use of a ubiquitous promoter would allow for expression in critical cell and tissue types.

**Figure 1.**
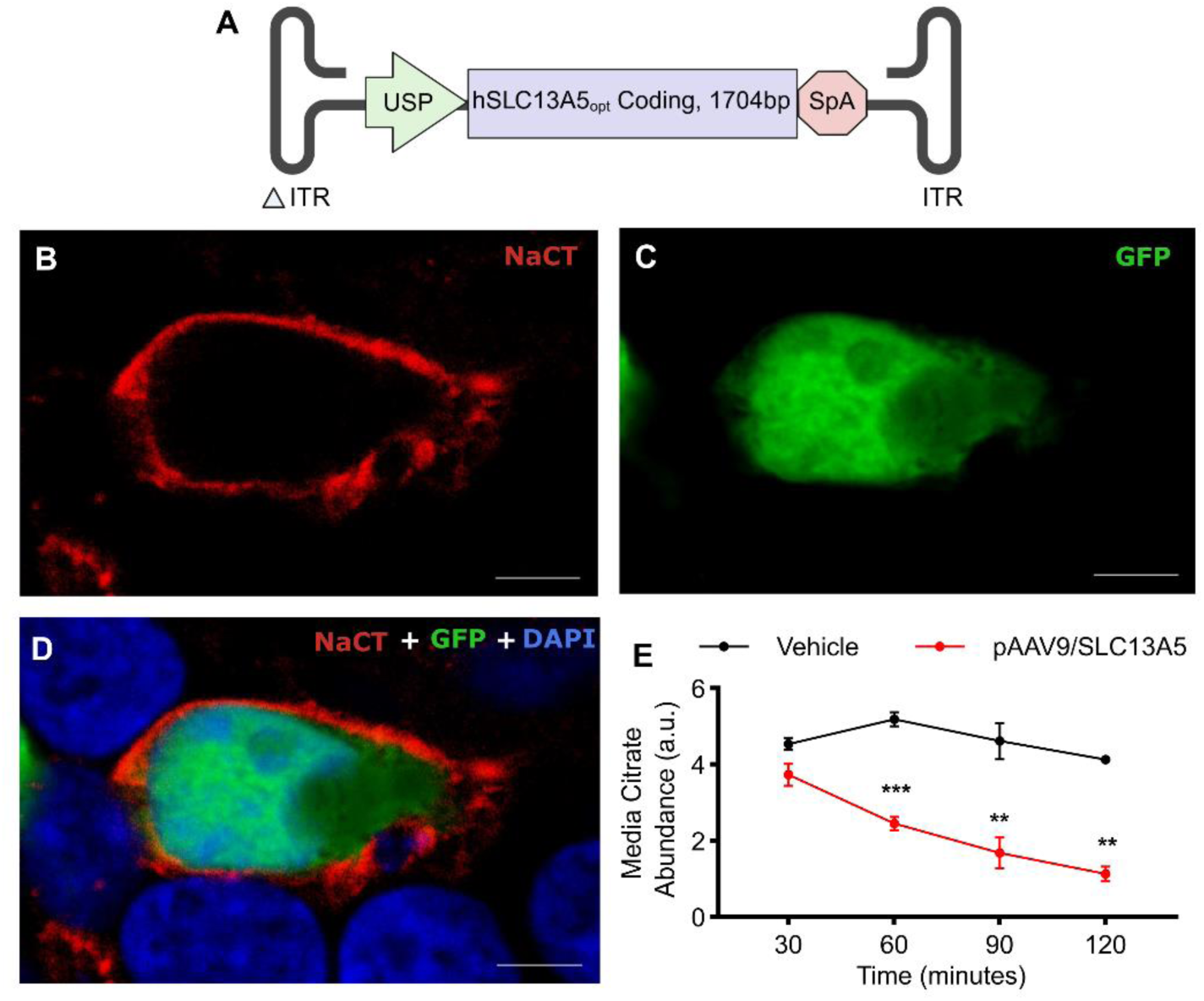
AAV/SLC13A5 vector design, expression and function in cells. **A.** Schematic of AAV9/SLC13A5 transgene. **B-D**. Representative image of a HEK293T cell co-transfected with a plasmid expressing eGFP and pAAV/SLC13A5 showed that 48 hours after transfection NaCT staining was in a punctate pattern around the outer edge of the cell and that eGFP fluorescence, which was detected homogenously in the cytosol, did not co-localize with NaCT (**B.** NaCT, **C.** GFP, **D.** merged NaCT, GFP and DAPI). *Scale bar* 5 µm**. E.** Uptake of citrate (1 µM) into HEK293T cells transfected with vehicle or pAAV9/SLC13A5. Citrate abundance in the media was assessed at 30, 60, 90 and 120 minutes after addition to wells via GC-MS. Two-way ANOVA, Sidak’s post-hoc analysis; **p<0.01, ***p<0.001. n=3 per treatment. a.u. = arbitrary units.

Expression of the AAV/SLC13A5 vector plasmid was assessed using immunocytochemistry as commercially available NaCT antibodies did not reliably identify a band via western blot analysis. HEK293T cells lack NaCT expression and have naturally low transport of extracellular citrate, thus providing an applicable assay to test the functional potential of the SLC13A5 gene therapy vector in human cells. To assess localization, HEK293T cells were transfected with our pAAV/SLC13A5 plasmid and a plasmid expressing eGFP protein. eGFP is a soluble protein that is diffusely located in the cytoplasm and should not overlap with plasma membrane proteins. After 48 hours, cells were fixed and immunocytochemistry performed with antibodies that recognize human NaCT and eGFP, respectively, to verify that NaCT localized to the plasma membrane (Figure 1B-D). To confirm vector expressed NaCT transports citrate into cells, HEK293T cells were transfected with pAAV9/SLC13A5 or vehicle for 48 hours. 1 µM of citric-acid was added to the cell culture media and then a time course of collected media samples was assessed for citrate using GC-MS. Over 2 hours, cells transfected with vehicle showed very low baseline citrate transport, while pAAV9/SLC13A5 treatment decreased extracellular citrate levels by ∼70% (Figure 1E), indicating increased citrate transport from the extracellular fluid into cells. Together these results demonstrate that pAAV9/SLC13A5 results in functional NaCT protein in human cells.

### AAV9/SLC13A5 treatment restored functional NaCT in Slc13a5 KO mice

To test the hypothesis that broad CNS and peripheral delivery of the human SLC13A5 gene using AAV9 can provide an effective treatment for SLC13A5 citrate transporter disorder, we utilized homozygous *Slc13a5* KO mice (10). Preclinical efficacy studies were performed in post-natal day 10 (P10) pups, comparing IT delivery of either 2e11 vector genome (vg; low dose) or 8e11 vg (high dose) and mice were followed up to ∼4 mo post-injection (Figure 2A). Vehicle treated WT and KO littermates served as controls. We first determined if AAV9/SLC13A5 treatment resulted in functional NaCT by measuring blood citrate levels. This is clinically relevant as patients have significantly elevated citrate levels in the blood and CSF. Similarly, *Slc13a5* KO mice had significantly elevated citrate levels (Figure 2B). Treatment with AAV9/SLC13A5 significantly decreased plasma citrate levels in KO mice in a dose-dependent manner as compared to control KO mice, where at 2 mo post-injection low dose treated mouse citrate levels were decreased to 87% ± 7.7% of WT levels and high dose treated mice were decreased to 65% ± 8.0% (Figure 2C). This decrease in blood citrate levels was well tolerated, with no adverse effects observed across all the treated animals as determined by body weight gain/maintenance, survival, and general activity (Supplemental Figure 1).

**Figure 2.**
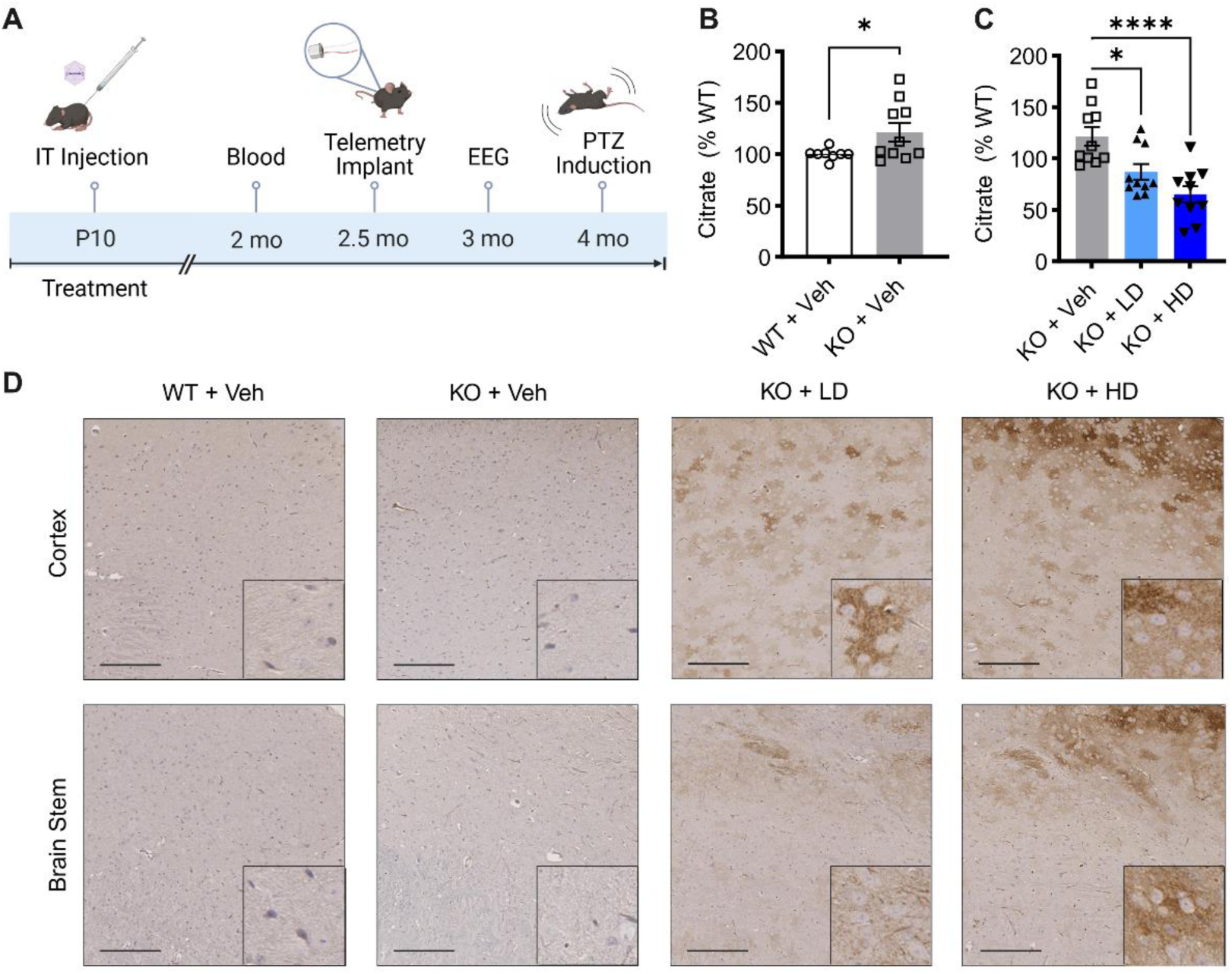
P10 IT administrations of AAV9/SLC13A5 gene therapy in *Slc13a5* KO mice resulted in robust NaCT in the brain and decreased blood citrate levels. (**A**) Schematic of the efficacy study in *Slc13a5* KO mice IT injected at P10 with vehicle, 2e11 vg (LD) or 8e11 vg (HD) of AAV9/SLC13A5. (**B-C**) GC-MS detection of citrate levels 2 mo post-treatment relative to WT controls. (**B**) Vehicle treated WT and *Slc13a5* KO mice. Student’s unpaired t-test, *p<0.05. (**C**) Vehicle, AAV9/SLC13A5 LD, and HD treated KO mice. One-way ANOVA with Dunnet’s Multiple Comparison’s test, *p<0.05, ***p<0.0001. n = 8 WT+Veh, 10 KO+Veh, 10 KO+LD, 10 KO+HD. (**D**) Representative images of mouse brain collected 4 mo post-treatment and stained with SLC13A5/NaCT antibody in the cortex (top) and the brain stem (bottom). *Scale bar* 200 μm. *Inset scale bar* 50 μm.

SLC13A5 transgene expression in the brain was confirmed by immunohistochemical analysis of NaCT protein in the brain. Staining in WT mice matched the staining in vehicle treated KO mice, indicating that the NaCT antibody used does not recognize mouse NaCT protein and that the staining observed is non-specific background signal (Figure 2D). In contrast, P10 AAV9/SLC13A5 treated mice showed dose-dependent, widespread expression of NaCT throughout the brain Figure 2D. Additionally, the morphology of the NaCT staining was consistent with that of a membrane protein as shown in Figure 2D insets, supporting that transgene expressed NaCT protein translocated to the plasma membrane.

### Seizure susceptibility was attenuated with AAV9/SLC13A5 treatment of Slc13a5 KO pups

Efficacy was tested, in part, by assessing epileptiform activity using wireless telemetry devices. Representative EEG traces demonstrate that ∼3 mo old *Slc13a5* KO mice had increased EEG abnormalities compared to WT mice (Figure 3A). Quantification of epileptic spike trains confirmed a small, non-significant increase in epileptiform discharges in control KO mice compared to WT mice (Figure 3B). In KO mice, treatment with AAV9/SLC13A5 decreased the number of spike trains in a dose dependent manner compared to vehicle treatment (Figure 3A-B).

**Figure 3.**
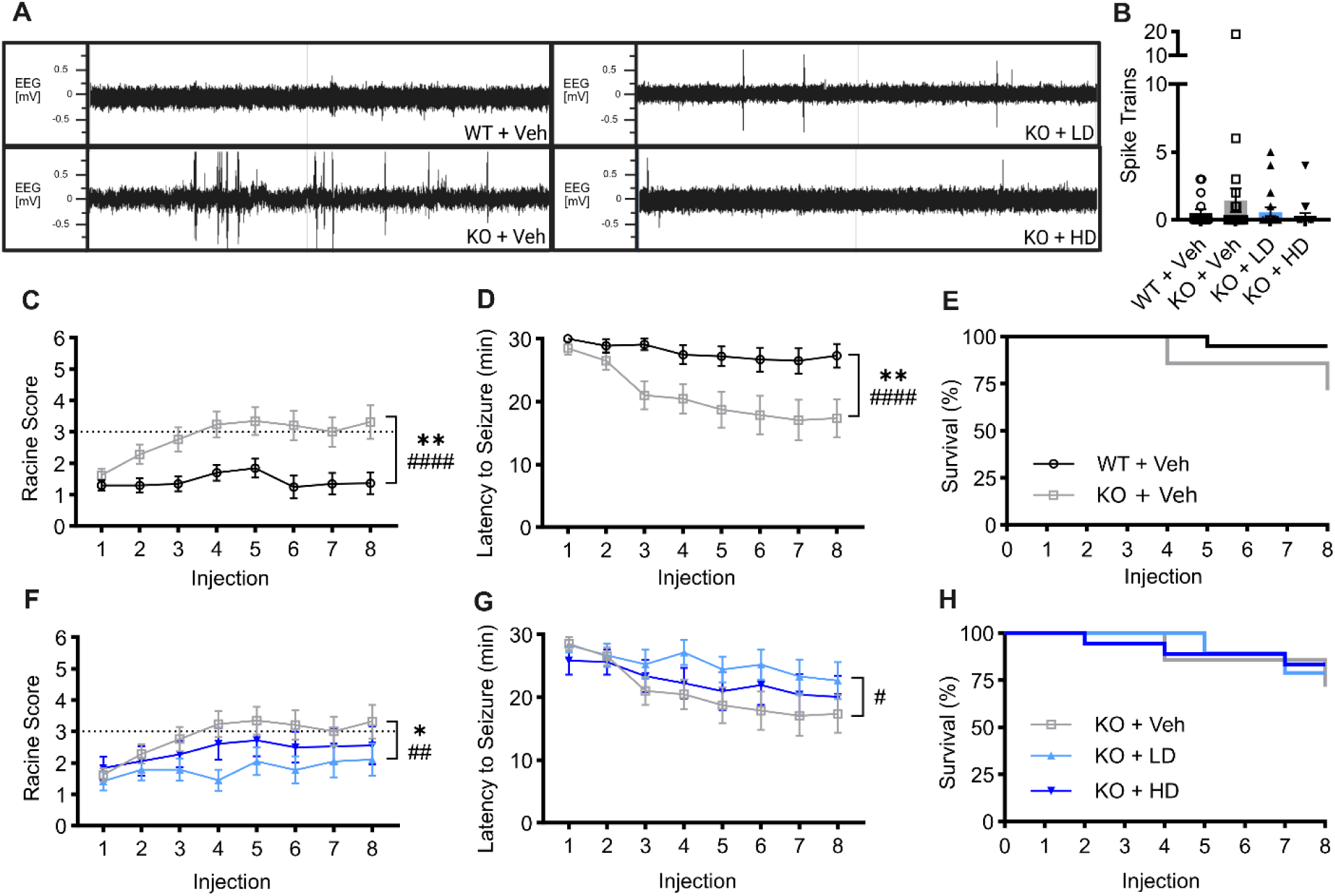
AAV9/SLC13A5 gene therapy rescued epilepsy in *Slc13a5* KO mice treated at P10. **(A-B)** Surface EEG was collected at 3 mo of age using a wireless telemetry implant in WT and KO mice IT injected at P10 with vehicle, 2e11 vg (LD) or 8e11 vg (HD) AAV9/SLC13A5. (**A**) Representative EEG traces over a 10 min period. (**B**) Total spike train counts over 2, 60-hour recordings. (**C-H**) Seizure susceptibility was assessed with administration of 30 mg/kg pentylenetetrazol (PTZ) every other day for 8 total injections. (**C-D**) Vehicle treated *Slc13a5* KO mice were significantly more susceptible to PTZ induced seizures compared to WT mice, exhibiting increased seizure severity (**C**) and decreased latency to seizure (**D**). (**E**) Survival of vehicle treated controls over the course of 8 injections. (**F-H**) Treatment with AAV9/SLC13A5 in KO mice reduced seizure severity (**F**) and increased seizure latency compared to KO+Veh mice **(G**). (**H**) Survival of PTZ study mice over the course of 8 injections. Repeated measures ANOVA as compared to KO+Veh group: genotype/treatment effect *p≤0.05, **p<0.01; genotype/treatment x PTZ injection effect ^#^p<0.05, ^##^p<0.01, ^####^p<0.0001. n = 20 WT+Veh, 21 KO+Veh, 19 KO+LD, 18 KO+HD. Data shown as Mean ± SEM.

*Slc13a5* KO mice have occasional sporadic seizures and are more susceptible to seizure induction (11). In young adult KO mice we did not observe sporadic seizures, so we tested seizure susceptibility using a chronic induction paradigm using a fixed, low dose (30mg/kg) of 8 injections of pentylenetetrazol (PTZ) (29). Seizure severity was scored using a modified Racine scale, where 0 to 2 represents none to partial/focal seizures and scores of 3 to 5 are generalized seizures and a maximal score of 6 represents prolonged tonic extensions of muscles that results in death (30). On average, WT mice failed to develop generalized seizures (score of 3 or greater) while KO mice had a significant increase in seizure severity that progressed to generalized seizures and coincided with a significant reduction in latency to seize (Figure 3C, D). KO mice treated with low-dose AAV9/SLC13A5 had significantly decreased seizure severity and increased the latency to seize with repeat PTZ injections as compared to vehicle treated KO mice (Figure 3F, G). KO mice treated with high-dose AAV9/SLC13A5 had similar Racine scores and latency to seize as low-dose treated KO mice and on average failed to develop generalized seizures. Additionally, 1/20 WT+Vehicle mice (5%), 6/21 KO+Vehicle mice (29%), 4/19 KO+LD (21%), and 3/18 KO+HD (18%) died by the 8^th^ injection (Figure 3E, H). Together, these results support that early gene therapy treatment reduces epileptiform activity and protects against seizure related death in a SLC13A5 mouse model.

### Adolescent AAV9/SLC13A5 treatment rescues EEG abnormalities and restores normal sleep

Previously, we reported that the vehicle treated young adult *Slc13a5* KO mice have increased activity during the light cycle, when mice typically sleep, decreased paradoxical sleep, and changes in absolute power spectral density as compared to the vehicle treated WT mice, indicating altered sleep architecture in KO mice (2). In treated KO mice, we found that during the dark cycle, when mice are typically active, vehicle and virus treated KO mice had similar levels of activity (Figure 4A). During the light cycle, the abnormally high activity in untreated KO mice was normalized with the high dose (Dunnett’s multiple comparisons test, p = 0.0061; Figure 4B), suggesting that treated KO mice spend more time asleep. Analysis of the specific sleep stages using the EEG and EMG data showed that vehicle and vector treated KO mice spent a similar percentage of time in active wake, quiet wake, and slow wave sleep stages (Figure 4C-E), while the low amount of time that vehicle treated KO mice spend in paradoxical sleep was increased with gene therapy in a dose-dependent manner (Dunnett’s multiple comparisons test, low dose p = 0.167, high dose p = 0.0492; Figure 4F).

**Figure 4.**
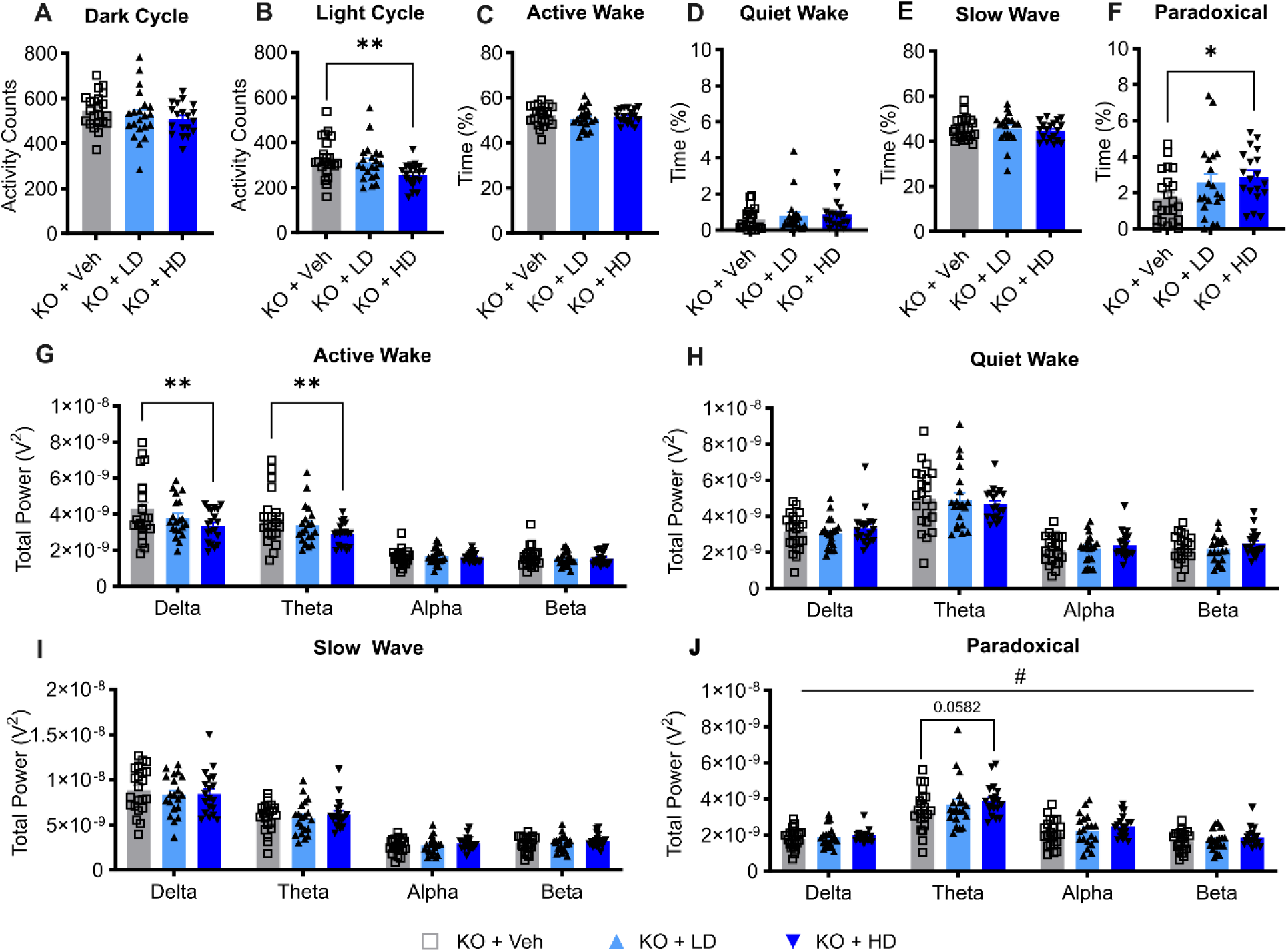
P10 IT delivery of AAV9/SLC13A5 rescued altered EEG sleep signatures in *Slc13a5* KO mice. (**A-B**) Overall activity counts of KO mice during the dark (**A**) and light (**B**) cycles. (**C-F**) Percentage of time spent in active wake (**C**), quiet wake (**D**), slow-wave sleep (**E**), and paradoxical sleep (**F**). One-way ANOVA with Dunnet’s Multiple Comparison’s test, *p<0.05, **p<0.01. (**G-J**) Power spectral density during active wake (**G**), quiet wake (**H**), slow-wave sleep (**I**), and paradoxical sleep (**J**), divided into delta, theta, alpha, and beta frequencies. Two-way ANOVA, treatment effect #p < 0.05; with Dunnett’s Multiple Comparison’s test, **p<0.01. n = 22 KO+Veh, 20 KO+LD, 19 KO+HD. Data shown as Mean ± SEM.

We then performed power spectral density (PSD) analysis across 0–30 Hz frequencies. Frequency bands were defined as delta (0.5–4 Hz), theta (4–8 Hz), alpha (8–12 Hz), and beta (12–30 Hz). Power band analysis of treated KO mice showed a dose-dependent decrease of slow wave activity (delta and theta waves) during the active wake stage (Dunnett’s multiple comparisons test, delta: low dose p = 0.1530, high dose p = 0.0024; theta: low dose p = 0.3327, high dose p = 0.0060). Previously, we reported that the absolute PSD during paradoxical sleep was significantly altered in vehicle treated KO mice as compared to vehicle treated WT mice, with theta and alpha power bands being significantly decreased in KO mice (2). In gene therapy treated mice, we found an increase of absolute PSDs during paradoxical sleep (two-way ANOVA, treatment effect (F (2, 232) = 3.574, p = 0.0296), with a dose-dependent increase of theta power waves (Dunnett’s multiple comparisons test, low dose: p = 0.3573, high dose: p = 0.0582), while alpha waves were similar amongst all treatment groups (Figure 4G-J). Together these results support that AAV9/SLC13A5 treatment in P10 KO mice achieves a dose-dependent rescue of sleep dysfunction.

### Gene therapy treatment resulted in a sustained reduction of CSF citrate levels

In addition to increased blood citrate levels, SLC13A5 mice also have elevated citrate levels in the CSF (8, 11). To determine if AAV9/SLC13A5 gene replacement therapy results in a sustained decrease of CSF citrate levels, a second cohort of mice was injected at P10 with 2e11 vg (low dose) and CSF was collected at 6-7 mo post-injection (Supplemental Figure 2A). Analysis of citrate via LC-MS confirmed that vehicle treated *Slc13a5* KO mice have significantly higher CSF citrate levels compared to vehicle treated WT mice (Supplemental Figure 2B). Gene therapy treatment decreased citrate levels to 81% ± 3.9% of WT levels (Supplemental Figure 2B). Together these results support that gene therapy treatment results in sustained, functional NaCT expression that decreases CSF citrate levels.

### ICM delivery AAV9/SLC13A5 in adult KO mice resulted in greater brain NaCT expression than IT delivery

Having found that gene therapy treatment in developing mice provided benefit, we then asked if treatment in adult mice, later in the disease course and after brain development is complete, is beneficial. To test this, young adult (3mo) *Slc13a5* KO and WT mice were treated with vehicle or AAV9/SLC13A5 and were followed for ∼6 mo post-injection (Figure 5A). Our prior work showed that IT lumbar puncture injection of AAV9 in adult mice results in significantly decreased brain transduction as compared to mice injected at P10 (31). Taking this into consideration, we assessed only the high dose (8e11 vg) in adult mice and compared IT delivery to intra-cisterna magna (ICM) delivery as we previously showed that ICM delivery in adult mice results in greater brain transduction as compared to IT delivery (32). Adult treatment was well tolerated, with no adverse effects observed across all the animals treated as determined by body weight gain/maintenance, survival, and general activity (Supplemental Figure 3).

**Figure 5.**
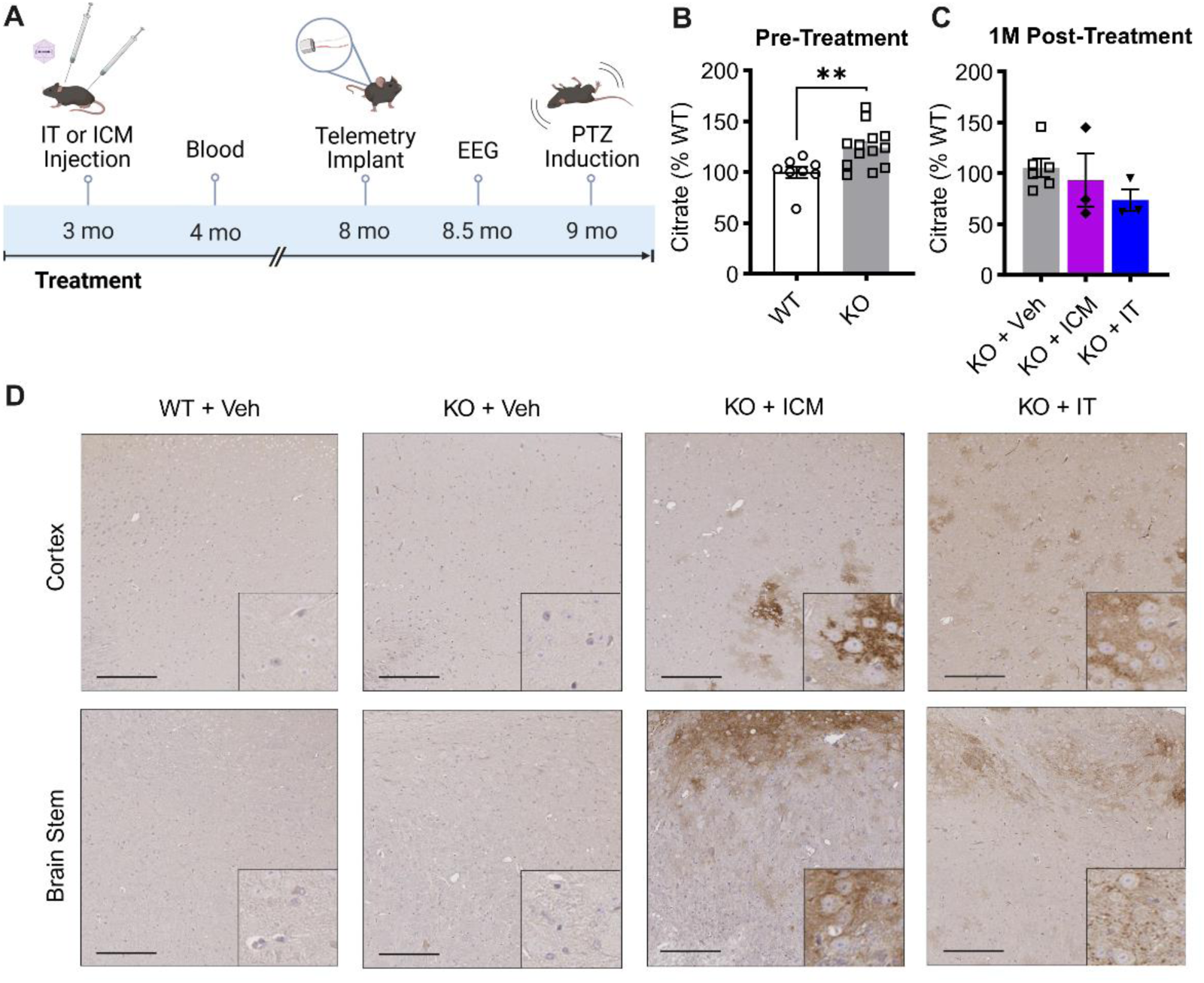
ICM delivery of AAV9/SLC13A5 in adult KO mice resulted in greater brain NaCT expression compared to IT delivery. **(A)** Schematic of the efficacy study in *Slc13a5* KO mice ICM or IT injected at 3 mo with vehicle or 8e11 vg AAV9/SLC13A5. (**B-C**) GC-MS analysis of plasma citrate, relative to WT+Veh controls. (**B**) Pre-treatment WT and KO controls. n = 8 WT, 13 KO. Student’s unpaired t-test, **p<0.01. (**C**) Vehicle, AAV9/SLC13A5 ICM, and IT treated KO mice 1 mo post treatment. n = 6 KO+Veh, 3 KO+ICM, 3 KO+IT. (**D**) Representative images of mouse brain collected 6 mo post-treatment and stained with SLC13A5/NaCT antibody in the cortex (top) and the brain stem (bottom). *Scale bar* 200 μm. *Inset scale bar* 50 μm.

Blood was assessed for citrate from a subset of WT and KO mice prior to AAV injection and 1 mo post injection. In agreement with our vehicle treated mice from our prior study (Figure 2), we found that untreated KO mice had significantly increased blood citrate levels compared to untreated WT mice (Figure 5B). Treatment with AAV9/SLC13A5 decreased plasma citrate levels in KO mice to 93.3% ± 26.1% of WT levels via ICM administration and to 73.7% ± 10.7% via the IT route (Figure 5C). We also assessed SLC13A5 transgene expression in the brain by staining for NaCT protein. In AAV9/SLC13A5 treated mice, there was widespread expression of the SLC13A5 protein throughout the brain, with higher expression achieved with ICM delivery as compared to IT delivery (Figure 5D). Compared to mice treated with the same dose and via the same route at P10 (Figure 2), KO IT injected at 3mo had relatively lower NaCT levels throughout the brain.

### AAV9/SLC13A5 treatment of adult KO mice rescued epileptic phenotypes

Mice were implanted with telemetry devices at ∼5 mo post-injection and EEG, EMG, and activity data was recorded over 2, 60-hour recording periods. In WT mice, EEG brain activity was normal while KO mice treated with vehicle had increased epileptiform activity (Figure 6A). ICM administration of AAV9/SLC13A5 normalized brain activity of KO mice to WT levels and to a lesser extent with IT administration (Figure 6A). Quantification of spike trains showed that compared to WT mice, KO mice had significantly increased number of spike trains and that treatment with AAV9/SLC13A5 resulted in a significant decrease in the number of spike trains with ICM administration and to a lesser extent with IT administration (Figure 6B).

**Figure 6.**
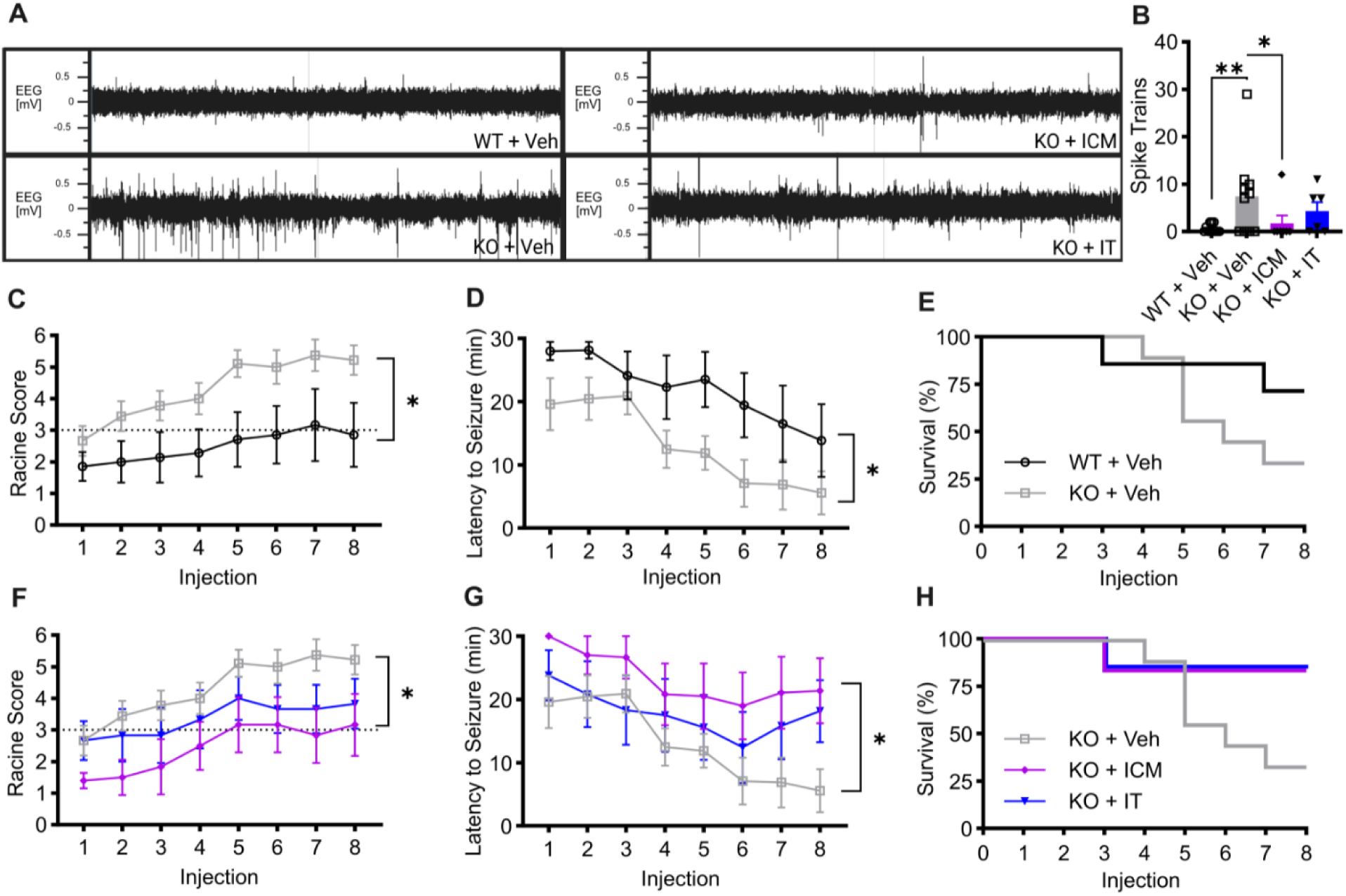
ICM delivery of AAV9/SLC13A5 had greater rescue of epileptic phenotypes compared to IT delivery in adult *SLC13A5* KO mice. (**A-B**) Surface EEG collected from WT and KO mice injected at 3 mo via ICM or IT with vehicle or 8e11 vg AAV9/SLC13A5. (**A**) Representative EEG traces over a 10 min period. (**B**) KO mice exhibited elevated spike train counts that were reduced with gene therapy. One-way ANOVA, *p<0.05, **p<0.01. (**C-H**) Seizure susceptibility was assessed with administration of 30 mg/kg PTZ every other day for 8 total injections. (**C-E**) Vehicle treated *Slc13a5* KO mice were significantly more susceptible to PTZ induced seizures compared to WT mice, exhibiting increased seizure severity (**C**), decreased latency to seizure (**D**), and increased incidence of seizure induced death (**E**). **(F-H**) Seizure susceptibility of AAV9/SLC13A5 treated mice were compared to KO+Veh mice. ICM delivery resulting in significant decreased seizure severity (**F**) and decreased latency to seizure (**G**). (**H**) ICM and IT delivery of AAV9/SLC13A5 reduced seizure induced death. Repeated measures ANOVA as compared to KO+Veh group: genotype/treatment effect *p≤0.05. n = 7 WT+Veh, 9 KO+Veh, 6 KO+ICM, 6 KO+IT. Data shown as Mean ± SEM.

Mice were then assessed for seizure susceptibility using our PTZ kindling paradigm described above. At 9 mo of age, WT mice had an average Racine Score of 2.9 ±1.0 by injection 8. In contrast, control KO mice were significantly more susceptible to seizure induction and had a mean Racine Score of 5.2 ±0.5 by injection 8 and had a significantly reduced latency to seize as compared to WT mice (Figure 6C, D). The increased seizure susceptibility resulted in early lethality in 2/8 WT+Vehicle mice (25%) and 6/9 KO+Vehicle mice (67%) (Figure 6E). Treatment of KO mice with AAV9/SLC13A5 decreased seizure severity with a significant improvement when delivered via the ICM route and to a lesser extent via the IT route as compared to vehicle treated KO mice (Figure 6F). By injection 8, KO+ICM mice and KO+IT mice had an average Racine Score of 3.2 ±1.0 and 3.8 ±0.8, respectively. AAV9/SLC13A5 treated KO mice also had increased latency to seizure onset as compared to KO control mice (Figure 6G). By decreasing seizure susceptibility, seizure related deaths were decreased in treated KO mice where only 1/6 KO+ICM (17%) and 1/6 KO+IT (17%) died by the 8^th^ injection (Figure 6H). Together, these results support that AAV9/SLC13A5 administration in young adult mice restored resistance to the onset of seizures and reduced seizure severity.

### Adult AAV9/SLC13A5 treatment did not alter general activity levels

Analysis of telemetry implant data showed that at 8 mo of age conscious, freely moving KO mice had similar activity levels within their home cages during the light and dark periods as compared to WT mice (Supplemental Figure 4). When treated mice were assessed, we found that injection of AAV9/SLC13A5 by either ICM or IT delivery route did not significantly alter the activity of older KO mice. (Supplemental Figure 4).

### Sleep dysfunction and abnormal electrophysiologic power spectrum were ameliorated in Slc13a5 KO mice when treated as adults

At 8 mo of age, KO mice spent significantly less time in the active wake phase and significantly more time in the slow wave sleep phase as compared to WT mice (Figure 7A-D). This supports that older adult *Slc13a5* KO mice have altered sleep physiology, although in contrast to young adult animals (2), sleep dysfunction was characterized by less time spent awake and more time spent asleep. When KO mice were treated with AAV9/SLC13A5, there was a trending increase in the amount of time spent in active wake and decreased time spent sleeping as compared to vehicle treated KO mice, with the greatest rescue seen in mice that were ICM treated versus IT treated (Figure 7E-H).

**Figure 7.**
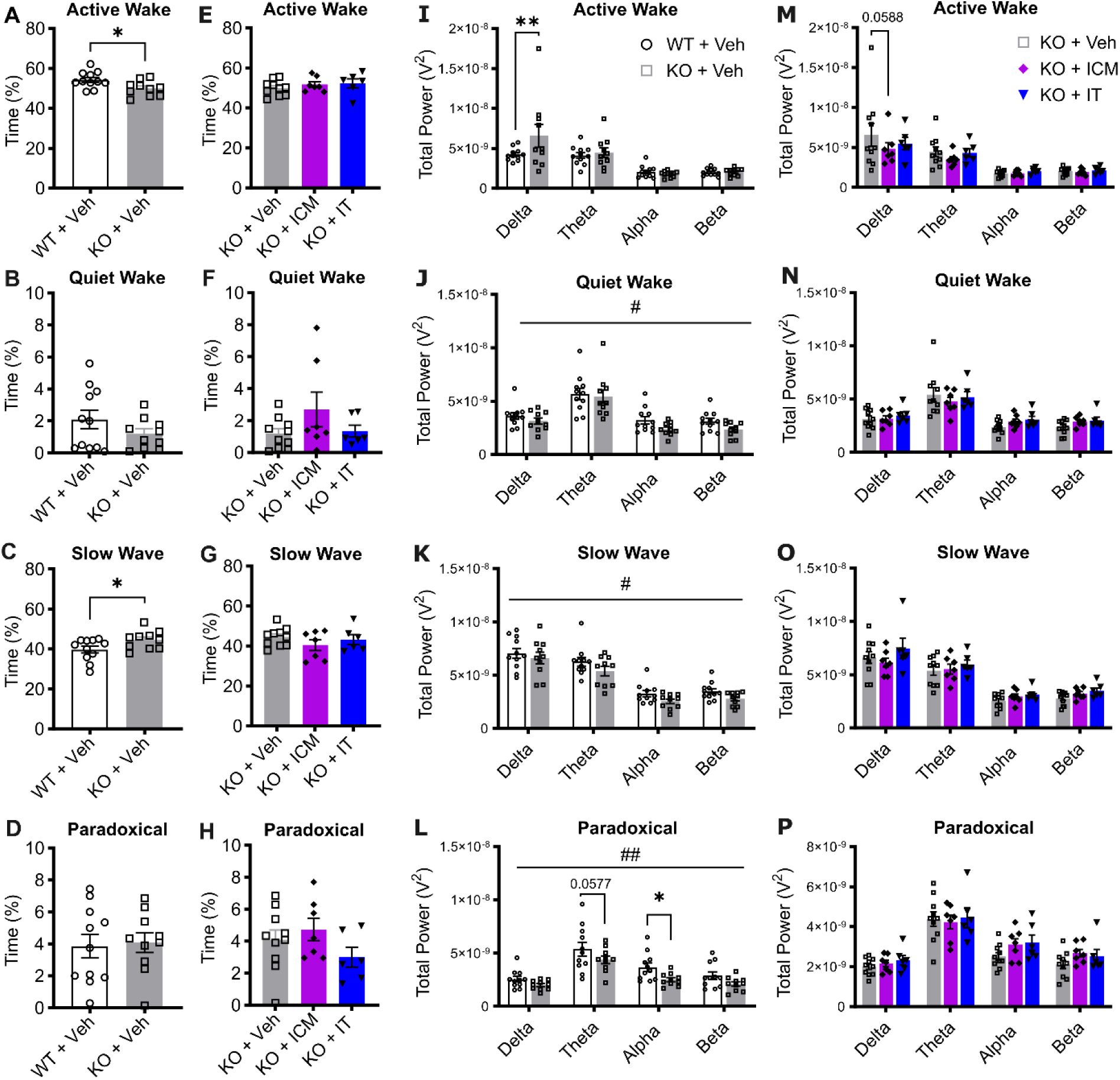
AAV9/SLC13A5 treatment in adult *Slc13a5* KO mice ameliorates sleep abnormalities. (**A-D**) Assessments of percentage of time spent in active wake (**A**), quiet wake (**B**), slow-wave sleep (**C**), and paradoxical sleep (**D**) in WT and KO vehicle treated mice. Student’s unpaired t-test, *p<0.05. (**E-H**) Assessments of percentage of time spent in active wake (**E**), quiet wake (**F**), slow-wave sleep (**G**), and paradoxical sleep (**H**) in vehicle, AAV9/SLC13A5 ICM, and IT treated KO mice. (**I-L**) Power spectral density during active wake (**I**) quiet wake (**J**), slow-wave sleep (**K**), and paradoxical sleep (**L**), divided into delta, theta, alpha, and beta frequencies in WT and KO Vehicle treated mice. Two-way ANOVA, genotype effect #p < 0.05, ##p<0.01; with Dunnett’s Multiple Comparison’s test, *p<0.05, **p<0.01. (**M-P**) Power spectral density during active wake (**M**) quiet wake (**N**), slow-wave sleep (**O**), and paradoxical sleep (**P**), divided into delta, theta, alpha, and beta frequencies in vehicle, AAV9/SLC13A5 ICM, and IT treated KO mice. n = 11 WT+Veh, 10 KO+Veh, 7 KO+ICM, 6 KO+IT. Data shown as Mean ± SEM.

The power spectra within the specific sleep/wake stages were then assessed. We found that delta waves significantly increased in KO mice during the active wake period as compared to WT mice (Figure 7I). In contrast, during quiet wake, slow wave sleep, and paradoxical sleep KO mice had an overall decrease of absolute PSD across all frequencies (including delta) as compared to WT mice (Figure 7J-L). In paradoxical sleep, theta, and alpha power bands were decreased in KO mice compared to WT mice (Figure 7L), which agrees with our prior power spectra analysis of 4 mo old KO mice (2). Treatment with AAV9/SLC13A5 decreased delta power during active wake and there was a general increase of the absolute spectral power during quiet wake, slow wave sleep, and paradoxical sleep (Figure 7M-P). Additionally, during paradoxical sleep, there was a trend of increased alpha activity in ICM and IT AAV9/SLC13A5 treated KO mice as compared to vehicle treated KO mice (Figure 7P). Together, these results support that older adult *Slc13a5* KO mice have sleep disturbances that result in increased slow wave power during wake periods, which can be ameliorated with AAV9/SLC13A5 treatment.

### Vector brain biodistribution was dose-dependent when administered in young mice and route dependent in adult mice

To assess vector distribution with the different ages of injection, routes of delivery, and doses, liver and brain (frontal cortex, brain stem, cerebellum) were assessed via dd-qPCR for SL13A5 transgene. Overall, *Slc13a5* KO animals dosed at P10 with the high dose had a significantly greater abundance of AAV9/SLC13A5 vg per μg of host genomic DNA when compared to those dosed with the low dose (Figure 8A). As expected for IT delivery of AAV9, the liver had a greater amount of vector than the brain (19, 32). KO animals dosed at 3 mo by ICM administration had a larger abundance of AAV9/SLC13A5 in the brain (cerebellum: 7.5×10^3^ ± 5.9×10^3^ vg/mouse genome) compared to animals dosed by IT administration (cerebellum: 1.2×10^3^ ± 7.4×10^2^ vg/mouse genome), while the amount of vector present in the liver was comparable (ICM: 1.5×10^4^ ± 5.2×10^3^ vg/mouse genome; IT: 1.4×10^4^ ± 2.8×10^3^ vg/mouse genome; Figure 8B). In agreement with our prior work in WT mice (31), IT delivery of AAV9 in KO mice at P10 resulted in markedly higher brain transduction (cerebellum: 15.7×10^3^ ± 3.1×10^3^ vg/mouse genome) as compared to KO mice IT injected with the same dose at 3 mo (cerebellum: 1.2×10^3^ ± 7.4×10^2^ vg/mouse genome). Together, the biodistribution data supports that CSF delivery of AAV9/SLC13A5 via IT lumbar puncture or ICM injection efficiently transduces target tissues important for SLC13A5 citrate transporter disorder, with greater distribution achieved with administration in younger animals versus older animals.

**Figure 8.**
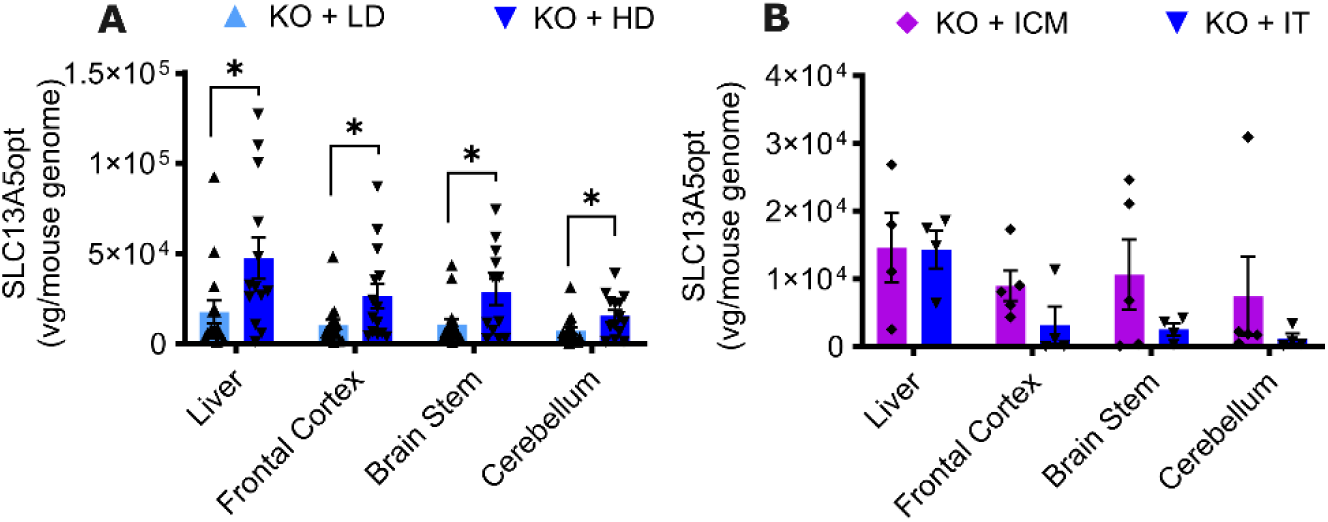
Biodistribution of AAV9/SLC13A5 following CSF delivery in *Slc13a5* KO mice. (**A-B**) Copies of AAV9/SLC13A5 per host genome in the liver, frontal cortex, brain stem, and cerebellum in mice dosed at P10 (**A**) and 3 mo (**B**). Student’s unpaired t=test, *p<0.05. n = 15 KO+LD, 14 KO+HD, 5 KO+ICM, 4 KO+IT. Data shown as Mean ± SEM.

## DISCUSSION

SLC13A5 citrate transporter disorder patients develop seizures within the first days of life and have lifelong epilepsy, neurocognitive impairments, a profound movement disorder, and limited verbal communication abilities that impact patient and family quality of life (4). Currently, there are no curative treatments that address the underlying disease cause, many patients require multiple anti-seizure medications simultaneously, and anti-seizure medications are not effective in all patients. Furthermore, patients require multiple additional medications, therapies, and support to address additional symptoms. We have developed a gene replacement approach for SLC13A5 deficiency that could have a profound impact on patients, potentially even into adulthood. Our preclinical studies showed that when packaged in AAV9 and delivered to *Slc13a5* KO pups or young adults, typical brain activity was restored, sleeping deficits were improved, and seizure resistance was increased. Results from these studies support that one-time CSF administration of AAV9/SLC13A5 is effective and sufficient in improving SLC13A5 deficiency-related dysfunctions.

SLC13A5 encoded NaCT is most highly expressed in liver, brain, bone, and reproductive organs, and is expressed in other organs to a lower extent (33-35). While NaCT plays a role in citrate uptake and energy metabolism in the liver, the impact of the liver phenotype on disease is unknown. The neurological phenotype is most prominent in SLC13A5 citrate transporter disorder (8, 36) and the greatest clinical benefit is likely to be achieved using strategies aimed at restoring SLC13A5 expression in the brain. We used IT or ICM administration of AAV9 as it results in higher transduction in the CNS and relatively lower transduction of peripheral tissues as compared to systemic delivery (18, 19). Analysis of vector biodistribution and expression in mice showed that P10 IT delivery resulted in widespread vector distribution and SLC13A5 expression throughout the brain. As expected, treatment with the high dose resulted in increased vector transduction and expression as compared to the low dose. Biodistribution analysis of tissues from adult-treated mice showed that IT and ICM administration of AAV9 resulted in similar liver transduction by both routes while ICM delivery resulted in greater brain expression as compared to IT injection. ICM delivery also provided the greatest rescue of epileptiform activity and restoration of seizure resistance, which supports brain NaCT deficiency as the main driver of disease. This also supports a lesser role for deficient liver NaCT expression for the neurological phenotypes we assessed. As AAV9 transduces both neurons and glia (27, 31, 32), it is unclear whether rescue of both cell types is necessary to ameliorate epileptiform activity or if restoring neuronal or glial NaCT function alone would be sufficient.

Administration of AAV9/SLC13A5 early in brain development provided therapeutic benefit to *Slc13a5* KO mice as assessed by citrate levels, brain activity, sleep behavior, and susceptibility to seizures and seizure-related death. Comparison of the low dose and high dose treated P10 mice showed benefit at the low dose that was even more beneficial with the high dose in most outcomes assessed, including EEG activity and sleep. Importantly, we also found that AAV9/SLC13A5 administration to adult mice, post neuronal development still had significant benefit. The potential for gene therapy to provide benefit when administered into early adult SLC13A5 patients is strengthened by the fact that although patients plateau in their development, there is no evidence of regression (1). SLC13A5 patients have structurally normal brains and while epileptiform activity is prominent, EEGs are typically not disorganized (5, 37). This indicates that brain architecture is largely in-tact in patients and suggests a wide age range for when gene therapy can be administered to restore functional NaCT and normalize brain activity.

Surprisingly in our PTZ kindling paradigm, low dose AAV9/SLC13A5 administered at P10 provided similar benefit as high dose. This may indicate a plateauing therapeutic effect with vector administration at P10. In transgenic mice, overexpression of SLC13A5 in a subpopulation of neurons (CAMKIIa+) beginning during embryonic development was reported to cause autistic-like and jumping behaviors, altered white matter integrity and synaptic plasticity, with aberrant synaptic structure and function (38). As S*lc13a5* gene expression in the rat cerebral cortex is low at birth and increases in the days following birth (39), the overexpression studies raise a concern that high levels of NaCT from vector expression during the prenatal or immediate post-neonatal period may impact proper brain development in rodents. While this could be contributing to the lack of further rescue of seizure susceptibility in the high dose P10 treated cohort, there was greater benefit with the high dose than low dose in all other metrics we measured including reducing epileptic discharges, normalizing brain waves, and ameliorating sleep deficits. Additionally, high dose AAV9/SLC13A5 treatment was well tolerated as mice had normal body weight, general activity, neurological function, and survival as compared to WT mice.

Epileptic phenotypes were notably different between younger and older adult mice, which can be reflective of alterations in inflammation, network function, and vulnerability to insult with aging. PTZ induced seizures in mice have been reported to worsen with age (40, 41), which was recapitulated within our findings, where WT mice tested at 4 mo of age had an average Racine score of 1.4 (Figure 3C) while WT mice tested at 9 mo of age had an average Racine score of 2.9 (Figure 6C). Further, we did not detect spontaneous seizures during the 60 hour recording periods in young adult KO mice while spontaneous seizures occurred in a few older adult vehicle treated KO mice, but not AAV treated mice. Similarly, the sleep phenotype varied with age. Three mo old KO mice had increased movement during the typical sleep period for rodents, and this coincided with decreased time spent during paradoxical sleep (2). In contrast, 8 mo old KO mice spent significantly less time in the active wake stage and more time in the slow wave sleep as compared to WT littermates, which could indicate that with aging abnormal excess sleepiness becomes a defining feature of SLC13A5 KO mice. Aging in mice and humans has also been shown to disrupt the ability to sustain sleep/wake states, particularly in wakefulness and the transition to NREM sleep (42, 43).

A consistent feature of young and older adult *Slc13a5* KO mice was lower theta and alpha waves during paradoxical sleep. AAV9/SLC13A5 treatment had a greater effect when treated at P10, where there was a significant decrease in hyperactivity during the sleep cycles (Figure 4B) and an increase in total paradoxical sleep (Figure 4F) and were similar to WT levels (2). AAV9/SLC13A5 treatment at 3mo of age had less effect in normalizing theta and alpha waves in KO mice (Figure 7O-P). This may be due to group sizes and being underpowered or indicates a neurodevelopmental feature of the disorder that does not rescue with adult vector administration. Interestingly, 8 mo old vehicle treated KO animals had significantly increased delta wave power during the active wake period as compared to WT littermates. Delta brain waves are typically a hallmark of sleep and should not be excessive during an awake state. Remarkably, gene therapy administration in adult animals was effective at decreasing delta activity during the active wake state, with greater normalization achieved with higher brain transduction (Figure 7M).

NaCT is responsible for transporting citrate from the extracellular fluid into the cell. In the absence of functional NaCT, citrate levels are distinctly increased in both the blood and CSF of patients and mice (7, 8, 10). While elevated extracellular citrate levels are a marker for impaired NaCT activity, they may not directly correlate with disease outcome (9). Our *Slc13a5* KO mice had significantly increased citrate in the blood and CSF, where citrate was ∼20% more in the blood and ∼50% more in the CSF as compared to WT levels. Treatment at P10 or at 3 mo significantly decreased both blood and CSF citrate levels, indicating functional NaCT activity. In treated mice, citrate levels decreased to as low as 65% of normal WT levels with the high vector dose at P10. It is unclear the clinical implications for low circulating citrate levels in the context of maintained citrate transport into cells. Circulating citrate that is not taken up by bones or tissues is filtered by the kidneys and excreted. Urinary citrate excretion by the kidneys has a critical role in maintaining acid-base homeostasis and in preventing calcium nephrolithiasis or formation of kidney stones (44). The reduction of circulating citrate was well tolerated in the *Slc13a5* KO mice and all tissues, including kidneys, were grossly normal at necropsy. Future studies, however, could assess citrate excretion following SLC13A5 gene therapy.

Compared to the clinical phenotype observed in patients with SLC13A5 citrate transporter disorder, *Slc13a5* KO mice have a relatively mild phenotype and lack significant motor and cognitive deficits that are important clinical features (45, 46) and a limitation of our studies was our inability to determine if gene therapy can improve motor and cognitive functions. These differences between mice and humans may indicate underlying biochemical (e.g., different K_M_) or physiological differences in NaCT function between species (45, 46). Currently, there is not an animal model that robustly models all aspects of the human disorder, and the KO mice were the best option available for testing our gene therapy. We were able to use *Slc13a5* KO mice to confirm NaCT protein expression following AAV9/SLC13A5 therapy, transporter function, and vector distribution within the brain and throughout the body. Additionally, changes in citrate levels, brain activity, sleep, and seizure susceptibility were used to determine if AAV9/SLC13A5 was beneficial and informed critical variables in therapy such as dosing, route of administration and age of treatment.

These results support that AAV9/SLC13A5 may be beneficial to patients with SLC13A5 citrate transporter disorder in restoring functional NaCT, restoring typical brain activity, normalizing sleep, and reducing epilepsy. Analysis of citrate in blood or CSF may serve as biochemical marker of functional NaCT being restored. Overall, our findings support further development and assessment of AAV9/SLC13A5 safety and efficacy.

## METHODS

### Sex as a biological variable

Our study examined male and female animals, and similar findings are reported for both sexes.

### Animals

C57BL/6J-*SLC13A5* KO mice were a gift from Dr. Rafael De Cabo and were generated as previously described (10). Mice were bred in UTSW animal barrier facilities under controlled environmental conditions, 12-hour light-dark cycles, and access to commercial 2916 chow and chlorinated reverse osmosis water ad libitum. Breeding pairs of either heterozygous males and females or homozygous males and heterozygous females were used to generate *SLC13A5* KO mice with litter, sex, and age matched controls.

### Viral vectors

We designed and developed the UsP-hSLC13A5opt-SpA plasmids (Figure 1A). Self-complementary AAV9 vector was produced using methods developed by the University of North Carolina (UNC) Vector Core facility. The purified AAV was dialyzed in PBS supplemented with 5% D-Sorbitol and an additional 212 mM NaCl (350 mM NaCl total). Vector was titered by qPCR and confirmed by polyacrylamide gel electrophoresis and silver stain. Two vector lots were used across studies. See Supplemental Figure 5 for the certificate of analysis for each lot.

### Virus Delivery

SLC13A5 homozygous KO and WT mice were treated at P10 or 3 mo. At P10, mice received an IT lumbar puncture injection of vehicle, 2e11 vg (low dose) or 8e11 vg (high dose) of AAV9/SLC13A5. At 3M, mice were injected with vehicle or 8e11 vg of AAV9/SLC13A5 via ICM or IT injection. Cage mates consisted of both vehicle and vector treated mice. On the day of injection, cage mates were randomly assigned to treatment groups, which were balanced for sex. For the IT injection procedure, mice were unanesthetized. Mice were held securely near the pelvic girdle and using a 30-gauge needle (Hamilton, Cat# 7803-07) with a Hamilton syringe (VWR, Cat# 89210-094) 5 μL of vector solution was slowly injected into the cerebrospinal fluid space between the L5 and L6 vertebrae. The needle was held in place for 2 seconds post-injection, then rotated, and withdrawn. For the ICM injection procedure, mice were anesthetized using an isoflurane vaporizer. Following induction of anesthesia and surgical prep, a small incision (5-10 mm) was made in the skin at the base of the skull. Lidocaine 2% (∼1-2 drops) was topically applied to the incision site. Using a 30-gauge needle (Hamilton, Cat# 7803-07) with a Hamilton syringe (VWR, Cat# 89210-094), 10 μL of vector solution was injected into the cisterna magna. The needle was then removed and the skin closed. Mouse body weight and survival were recorded from date of injection. Animals were then followed up to 6 mo post-injection and assessed for brain electrophysiologic activity, seizure susceptibility, blood citrate levels, vector biodistribution, and NaCT expression. See Supplemental Table 1 for injection cohorts, study numbers, and treatments.

### Immunocytochemistry

HEK293T cells were maintained in DMEM with 10% Fetal Bovine Serum, 1% glutamax, and 1% Pen/Strep. Sterile 12 mm coverslips were coated with 50ug/ml of Poly-D-Lysine in borate buffer. HEK-293T cells were plated on coverslips in a 24 well culture plate at a density of 5.0 × 10^4^ cells per well. 24hrs after plating, cells were transfected with 0.5ug GFP plasmid alone or in combination with 0.5ug plasmid AAV/SLC13A5opt using Lipofectamine 2000 in Opti-MEM. 48hrs post-transfection, coverslips were fixed using 4% paraformaldehyde for 30 min and permeabilized with 0.2% Triton in 1x PBS for 10 minutes. Coverslips were stained with SLC13A5 antibody overnight at 4°C (1:500; H00284111-M06, Abnova) and Alexafluor 594 secondary antibody (1:1000) for 1 hour at room temperature. Coverslips were counterstained with DAPI (500 ng/mL) and mounted with Aqua Poly-mount (18606, Polysciences). Slides were imaged using confocal microscopy (Zeiss LSM 780).

### Citrate Analysis

For citrate uptake analysis, HEK293T cells were maintained and plated in DMEM with 10% Fetal Bovine Serum, 1% glutamax, and 1% Pen/Strep. 24 hours after plating, cells were transfected with pAAV9/SLC13A5 or vehicle. After 48 hours, 1 µM of citric-acid was added to the cell culture media and samples of cell culture media were then collected at 30, 60, 90, and 120 minutes after the addition of citrate. Citrate in media was then quantified using GC-MS.

For citrate analysis, blood samples were collected from the facial vein using 4mm lancets (Fisher, NC9922361) into tubes containing 0.5M EDTA to be processed for plasma. Samples were centrifuged at 1500g at 4°C for 12 minutes and then snap frozen in liquid nitrogen. Blood was drawn from P10 cohort animals 1 mo post-injection. Blood was drawn from mice in the 3 mo cohort prior treatment and 1 mo post-injection. For terminal CSF harvests, at 6-7 mo post-injection, animals were deeply anesthetized by isoflurane and were secured under a stereotaxic microscope. A sagittal incision was made in the skin along the back of the skull, carefully separating the subcutaneous tissue and neck muscle through the midline to reveal the cisterna magna. Harvests were completed by puncturing the cisterna magna using a pulled capillary (Fisher, NC9544044), collecting between 2-7 uL per animal, ensuring no blood contamination within the sample. Samples were immediately snap frozen in liquid nitrogen. Citrate levels in plasma and CSF citrate were quantified using GC-MS analysis.

### Telemetry devices, recordings, and analysis

Animals were subcutaneously implanted with HD-X02 (Data Systems International, DSI) telemetry probes. Dural EEG leads were placed using the coordinates LH: AP +1.0, ML −1.5, RH: AP −2.0, ML +2.0 with EMG being placed in the trapezius muscles in the back of the neck. Following ∼10 days of recovery from surgery, EEG and EMG recordings were acquired over the course of two, 60-hour sessions (3 Dark Cycles, 2 Light Cycles) approximately two weeks apart. All the recordings took place singly housed in the animal’s home cage with food and water ad libitum. Ponemah (Data Systems International, v6.51) was used to acquire data, and Neuroscore (Data Systems International, v.3.4.0) was used to analyze data. Epileptic spike trains were assessed using a 10 Hz high pass filter with the automatic spike analysis protocol. Activity was automatically determined by implant movement within the cage. These counts were separated into activity during the dark cycles and activity during the light cycles, summed for two (light) or three (dark) cycles per recording period. Sleep architecture was categorized into active wake, quiet wake, slow wave sleep, or paradoxical sleep based on delta and theta power, muscle tone, and movement using the integrated Rodent Sleep Scoring program. The percent time spent in each sleep stage was averaged over the two recording sessions. Power Spectral Density Absolute was automatically quantified by the software for four frequency bands: delta (0.5–4 Hz), theta (4–8 Hz), alpha (8–12 Hz), and beta (12–30 Hz). Power within the sleep stages was averaged over the two recording sessions. Mice that died prior to recording did not have EEG data collected (P10 treatment: n = 5 WT+Veh, 4 KO+Veh, 5 KO+LD, 3 KO+HD; 3 mo treatment: n = 2 KO+Veh, 1 KO+IT). Additionally, due to automatic light malfunction during a recording session, EEG data was excluded from 3 WT+Veh P10 treated mice.

### Seizure Induction

Seizure susceptibility was assessed using a kindling PTZ induction paradigm. Mice received a 30 mg/kg intraperitoneal injection of PTZ (18682, Cayman Chemicals) dissolved in a sterile 0.9% saline solution every other day for a course of eight injections (29). Because of telemetry implants, mice were lightly anesthetized using isoflurane for PTZ administration. Animals were monitored for 30 minutes following PTZ injection and seizure severity was scored using a modified Racine scale (11): Stage 0 (no response); Stage 1 (immobilization); Stage 2 (myoclonic jerks); Stage 3 (clonic seizures); Stage 4 (tonic seizures with rearing); Stage 5 (generalized tonic-clonic seizures); Stage 6 (seizure induced death). For seizure latency, time to Stage 3 was recorded. Mice that died on day 1 of injections (P10 treatment: n = 1 KO+Veh, 1 KO+LD, 1 KO+HD; 3mo treatment: n = 4 WT+Veh, 1 KO+Veh, 1 KO+ICM) were excluded from PTZ analysis as this indicates a hypersensitivity to PTZ.

### Necropsy

At study end, animals were anesthetized via avertin overdose (0.05mL/g of a 2.5% solution), a terminal blood draw was taken *via* cardiac puncture, and tissues were collected after perfusion with PBS containing 1ug/mL heparin. Tissues collected were either snap frozen in liquid nitrogen for vector distribution or fixed in 10% NBF for histological analysis. Frozen tissues included liver in addition to brain hemisphere sub-dissected into frontal cortex, brainstem, and cerebellum.

### Vector biodistribution

Total genomic DNA was purified from frozen tissue samples using a QIACube HT (Qiagen) and following the 96 DNA QIACube HT Kit protocol. For vector expression, RNA was extracted in QIAzol lysis reagent and chloroform, and purified using a QIAcube HT. Extracted RNA samples were treated with DNAse I (M0303S, NEB), and cDNA synthesis performed using a Transcriptor First Strand cDNA Synthesis Kit (4896866001, Roche). qPCR analysis was performed by Northern Biomolecular Services.

### SLC13A5 Immunohistochemistry

Fixed brains were submitted to the UTSW Histopathology Core for processing, paraffin embedding and sectioning. Slides were generated with 5 μM thick serial sections. Slides were deparaffinized using standard methods and antigen retrieval was performed with antigen unmasking solution (Vector H-3300). The primary antibody used was mouse monoclonal anti-SLC13A5, clone 2G4 (Abnova H00284111-M06, 1:400). The secondary antibody used was biotinylated anti-rabbit (Vector BA-9200; 1:400). Slides were counterstained with Mayer’s hematoxylin and mounted using Poly-Mount. Slides images were captured using a NanoZoomer and images exported with the ImageScope Software (Leica Biosystems Inc., Buffalo Grove, IL).

### Statistics

Log-rank (Mantel-Cox) test was used to compare survival curves. Body weight was analyzed using a repeat measures ANOVA, with factors treatment or days-post injection, and followed with Dunnett’s multiple comparison test. PTZ was analyzed using a repeated measured ANOVA with factors for injection and genotype or treatment and followed with Dunnett’s multiple comparison test. Spike train counts, sleep architecture, and citrate were analyzed using a one-way ANOVA. Biodistribution was analyzed using a student’s unpaired t-test. For all comparisons, statistical significance was set at p ≤ 0.05. Data were analyzed and graphed using GraphPad Prism software (v. 10.4.2). Graphics were created with Biorender.com.

### Study approval

All animal experiments were reviewed and approved by the UTSW Institutional Animal Care and Use Committee.

## Conflict of Interest

RMB is an inventor on the SLC13A5 vector design (US 2023.0285595) and has received income related to that invention.

## Data Availability

All data associated with this study are available in the main text or the supplemental materials, and additional study details are available from the authors upon request. We previously reported analysis of sleep architecture and power spectra from WT+Veh and KO+Veh mice in the P10 cohort in Adams et al. (2). In the current manuscript, new analysis of KO+Veh, KO+LD, and KO+HD mice from the P10 cohort is reported. Any plasmids, vectors, or animal models used in this study are available upon request through a material transfer agreement. Values for all data points in graphs are reported in the Supporting Data Values file.

## AUTHOR CONTRIBUTIONS

RMB conceptualized the study. LB, MS, RMA, IG, KK, SH, and ME performed experiments. LB, RMA, and RMB analyzed data. ML helped with statistical analysis. RMB supervised all activities of the study. LB, RMA, and RMB wrote the original manuscript draft. All authors reviewed the manuscript.

## ACKNOWLEDGEMENTS

We thank Dr. Rafael De Cabo at the National Institute on Aging for providing the *Slc13a5* KO mouse line. We thank Dr. Xin Chen for his technical help with IT injections in adult mice. We thank the UTSW Histopathology Core for histological services and the UTSW Whole Brain Microscopy Facility (RRID:SCR_017949) for assistance with slide scanning. We thank the UTSW Neuro-models Facility for assistance with telemetry implant surgery and EEG recording (RRID:SCR_022529). We additionally thank the UTSW CRI Metabolomics facility for GC/LC-MS. We thank Drs. Brenda Porter and Tanya Brown for their review of this manuscript. The study was funded by the TESS Research Foundation and Taysha Gene Therapies.

