## Supplemental Figures for "AAV-mediated gene therapy for SLC13A5 Citrate Transporter Disorder rescues epileptic and metabolic phenotypes"

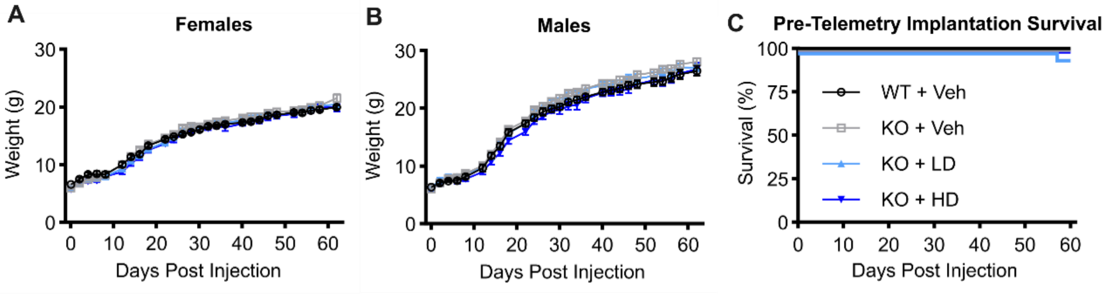


**Supplemental Figure 1: P10 IT treatment of AAV9/SLC13A5 resulted in normal weight gain and survival in KO mice.** WT and KO Slc13a5 littermates were treated with vehicle, 2e11 vg (LD) or 8e11 vg (HD) at post-natal day 10. (**A-B**) Weight gain in females (**A**) and males (**B**) following treatment up to 60 days post injection. (**C**) Survival following treatment until telemetry implant surgery, 60 days post injection. n = 12F/13M WT+Veh, 14F/12M KO+Veh, 13F/12M KO+LD, 11F/11M KO+HD. Data shown as Mean ± SEM.

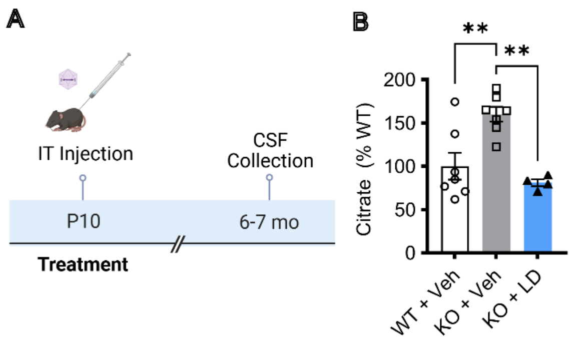


**Supplemental Figure 2: P10 IT administration of AAV9/SLC13A5 decreased CSF citrate in KO mice**. (**A**) Schematic of the efficacy study in Slc13a5 KO mice IT injected at P10 with vehicle, or 2e11 vg (LD). (**B)**. LC-MS analysis of citrate in CSF at 6-7 mo post-injection, relative to vehicle treated WT mice. One-way ANOVA as compared to KO+Veh with Dunnett's Multiple Comparison's test, **$p<0.01$. n=7 WT+Veh, 7 KO+Veh, 4 KO+LD. Data shown as Mean ± SEM.

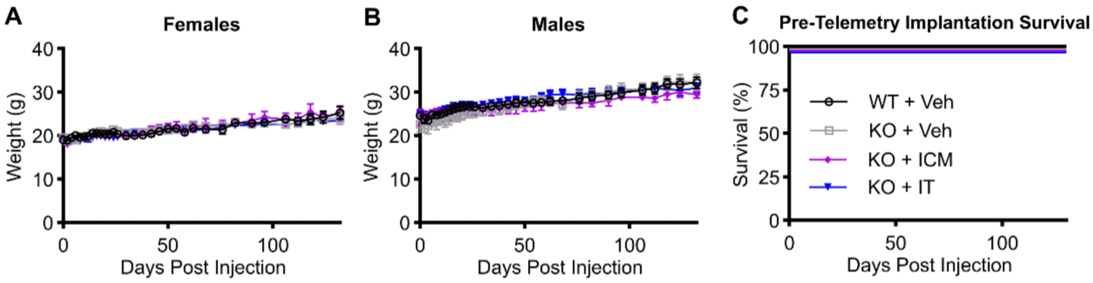


**Supplemental Figure 3: ICM and IT delivery of AAV9/SLC13A5 at 3 mo resulted in normal weight gain and survival in KO mice.** WT and KO Slc13a5 littermates were treated with vehicle or 8e11 vg via ICM or IT injection at 3 mo of age. (**A-B**) Weight gain in females (**A**) and males (**B**) following treatment up to 130 days post injection. (**C**) Survival following treatment until telemetry implant surgery, 60 days post injection. n = 5F/6M WT+Veh, 6F/6M KO+Veh, 3F/4M KO+ICM, 4F/3M KO+IT. Data shown as Mean ± SEM.

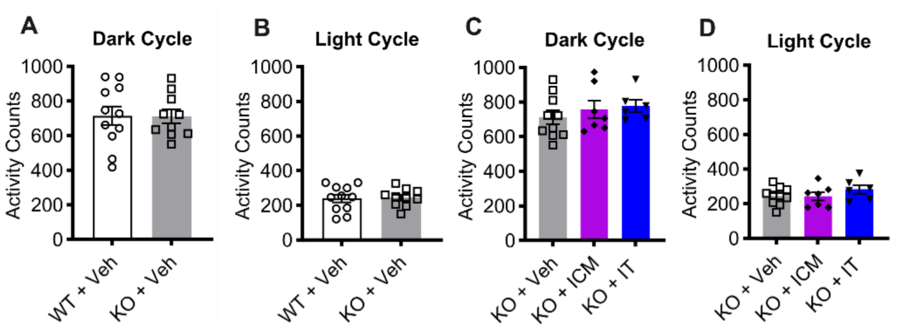


**Supplemental Figure 4: Slc13a5 KO mice had normal activity levels at 8 mo of age** (**A-B**) Overall activity counts of KO mice during the dark (**A**) and light (**B**) cycles at baseline. (**C-D**) AAV9/SLC13A5 treatment did not affect overall activity during the dark (**C**) or light (**D**). n = 11 WT+Veh, 10 KO+Veh, 7 KO+ICM, 6 KO+IT. Data shown as Mean ± SEM.

### Quality Control Summary

| Lot # | LAV55-conc |
|---|---|

**Test by qPCR**

| Test # | Titer, vg/mL | Analyst | Date | File |
|---|---|---|---|---|
| 1 | 1.55E+14 | PZ | 04/04/2018 | 20180404-1428-ghbh-pz |

**PAGE analysis**

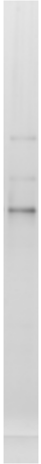

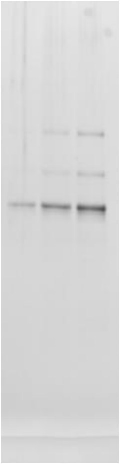

Loaded 5.00E+09 vg    4980E std  2e9vg  5e9vg  1e10vg
Calculated 4.30E+09 vg

| | |
|---|---|
| Analyst | Ping Zhang |
| Date | 04/02/2018 |
| Reference # | 20180402-silver |

**SEM**

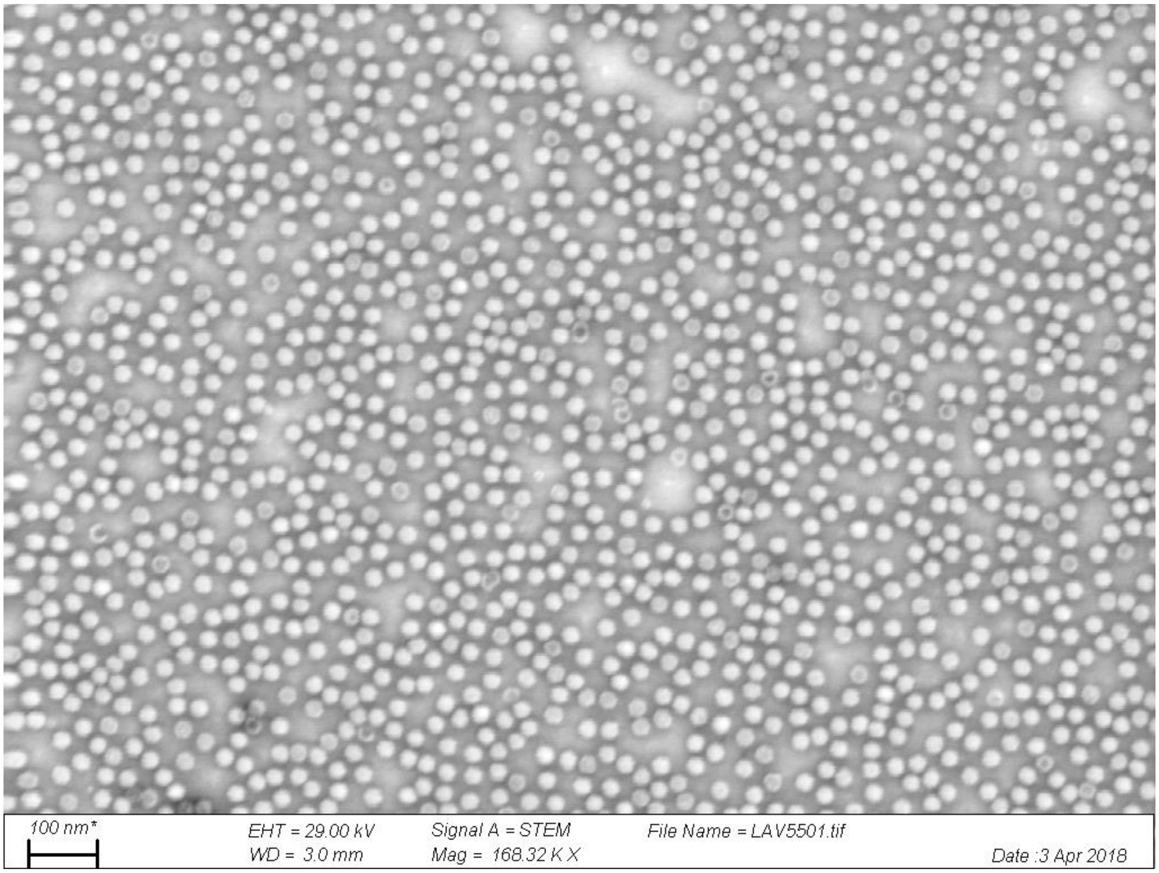


90% full

| | |
|---|---|
| Analyst | Ping Zhang |
| Date | 04/03/2018 |
| Reference # | 20180403-LAV55-01 |

### Quality Control Summary

| Lot # | LAV131 | Name | SLC13A5 |
|---|---|---|---|

**Test by qPCR**

| Test # | Titer, vg/mL | Analyst | Date | File |
|---|---|---|---|---|
| 1 | 1.38E+14 | PZ | 07/08/2020 | 20200708-1351-ghbh-pz |

**PAGE analysis**

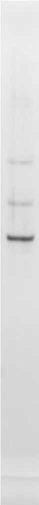

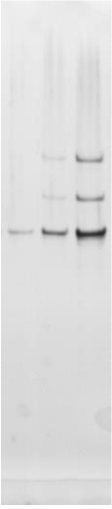

Loaded 5.00E+09 vg 4980E std 2e9vg 5e9vg 1e10vg

Calculated 5.10E+09 vg

| | |
|---|---|
| Analyst | Ping Zhang |
| Date | 06/29/2020 |
| Reference # | 20200629-silver |

**SEM**

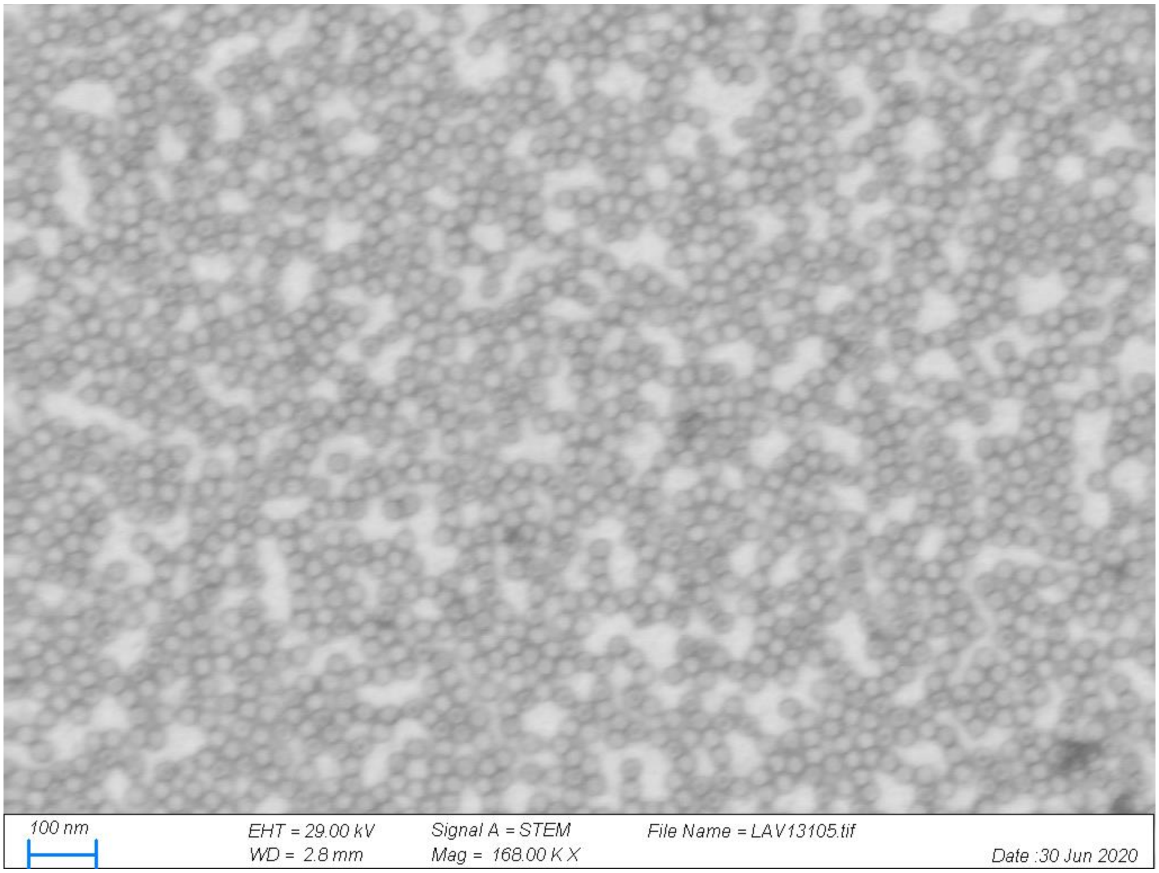


92% full

| | |
|---|---|
| Analyst | Ping Zhang |
| Date | 06/30/2020 |
| Reference # | 20200630-LAV131-05 |

**Supplemental Figure 5:** Certification of Analysis for AAV9/SLC13A5 lots.

#### Supplemental Table 1: Study animal numbers

| Study | Group | Treatment | EEG Recording | PTZ Induction | Biodistribution |
|---|---|---|---|---|---|
| **P10 Treatment** | WT + Veh | 25 (12F/13M) | 17 (9F/8M) | 20 (10F/10M) | - |
| | KO + Veh | 26 (14F/12M) | 22 (11F/11M) | 21 (10F/11M) | - |
| | KO + LD | 25 (13F/12M) | 20 (10F/10M) | 19 (10F/9M) | 15 (8F/7M) |
| | KO + HD | 22 (13F/12M) | 19 (9F/10M) | 18 (9F/10M) | 14 (7F/7M) |
| **3 mo Treatment** | WT + Veh | 11 (5F/6M) | 11 (5F/6M) | 7 (5F/2M) | - |
| | KO + Veh | 12 (6F/6M) | 10 (4F/6M) | 9 (4F/6M) | - |
| | KO + ICM | 7 (3F/4M) | 7 (3F/4M) | 6 (3F/3M) | 5 (3F/2M) |
| | KO + IT | 7(4F/3M) | 6 (4F/2M) | 6 (4F/2M) | 4 (3F/1M) |
